# Elevated CO₂ enhances tomato tolerance to *Botrytis cinerea* through transcriptional and metabolic defence reprogramming

**DOI:** 10.64898/2026.07.22.740109

**Authors:** Francesca Baistrocchi, Marta Orero-Bayo, Lili Yu, Victoria Pastor, Rosa Sanchez-Lucas, Antoni Garcia-Molina

**Affiliations:** Centre for Research in Agricultural Genomics (CRAG) CSIC-IRTA-UAB-UB, C/Vall Moronta, Edifici CRAG, 08193 Bellaterra (Cerdanyola del Vallès), Barcelona, Spain; Department of Biology, Biochemistry, and Natural Sciences, School of Technology and Experimental Sciences, Universitat Jaume I, 12006 Castelló de la Plana, Spain; Department of Biochemistry and Molecular Biology, University of Cordoba (UCO), Campus de Excelencia Internacional A3 (CeiA3), E-14014 Cordoba, Spain

## Abstract

Atmospheric CO₂ concentration is projected to rise substantially over the coming decades, yet its impact on the molecular mechanisms governing plant immunity remains poorly understood. Here, we investigated how elevated CO₂ (eCO₂; 650 ppm) combined with increased temperature (+5 °C) influences tomato responses to *Botrytis cinerea* through integrated phenotypic, metabolomic, transcriptomic, and gene regulatory network (GRN) analyses across eight cultivars. Although cultivars displayed contrasting susceptibility under ambient conditions, eCO₂ consistently enhanced tolerance across all genetic backgrounds. Multi-omics analyses revealed a partial uncoupling between transcriptional and metabolic responses during infection, with repression of photosynthesis- and carbon metabolism-related genes contrasting with the accumulation of carbon- and amino acid-derived metabolites. Under eCO₂, this metabolic disruption was attenuated, preserving metabolic homeostasis during infection. GRN reconstruction identified a conserved WRKY–ERF regulatory module underlying the growth–defence trade-off, while functional perturbation demonstrated that its contribution to resistance depends on both genotype and environmental context, highlighting the importance of basal defence mechanisms. Targeted metabolomics further revealed that eCO₂ promotes a metabolically primed state characterized by reinforcement of structural and chemical defence barriers rather than stronger activation of inducible immune responses. Together, our findings show that enhanced tolerance under eCO₂ emerges from coordinated reorganization across regulatory and metabolic networks, providing a systems-level framework for understanding plant immunity and improving crop resilience under future climate scenarios.

## Introduction

Elevated atmospheric CO2 (eCO2) is a defining feature of ongoing climate change and a major driver of plant physiological responses. Increased CO2 availability typically enhances photosynthetic carbon assimilation and promotes growth, leading to substantial changes in primary metabolism and biomass accumulation (Thompson *et al*., 2017; Kaiser *et al*., 2017). While the effects of eCO2 on plant productivity are well documented for different crop species (Dong *et al*., 2018; Lanoue *et al*., 2018; Doddrell *et al*., 2023; Luo *et al*., 2026), they pose an unprecedent interplay with responses to stressors.

Plant immunity relies on physical barriers (trichomes, stomata, epidermis and cell wall reinforcement) and molecular programs to sense broad spectrum of microbial epitopes (Pattern Triggered Immunity) and to neutralise pathogen’s virulence effectors (Effector Triggered Immunity) (Couto and Zipfel, 2016; Cui *et al*., 2015). In a systems-wide perspective, immunity consists of sophisticated interplays among multiple and diverse components that act co- ordinately to counteract the progression of the infection (Jones and Dangl, 2006). Hence, the integration of the different episodes of the infection triggers reconfigurations of plant physiology and metabolism (macroscopic level) orchestrated by fine-tuned rewiring in homeostatic networks (microscopic level), mediating transitions from the basal to distinct immune states. However, the energetic cost of maintaining immune states can trade off plant fitness and therefore the intensity and amplitude of defence responses are strictly tailored and gradually implemented depending on plant resources (Jones and Dangl, 2006; Bostock *et al*., 2014). Indeed, prolonged episodes of infection surpassing plant’s capacity to respond leads to a hypersensitive response triggering local programmed cell death to avoid the infection to spread (Bostock *et al*., 2014; Jones and Dangl, 2006). This growth-defence balance has traditionally been viewed as a trade-off, where increased investment in growth limits the resources available for immune responses. However, growing evidence suggests that this relationship is more flexible than previously assumed and may be strongly influenced by environmental conditions. Increased carbon availability under eCO2 may provide additional resources to sustain or even enhance defence responses, depending on how metabolic fluxes are reprogrammed (Bazinet *et al*., 2022; Li and Ahammed, 2023; Sanchez-Lucas *et al*., 2023; Smith and Luna, 2023). Nonetheless, the mechanisms by which elevated CO₂ influences the coordination between growth and defence remain poorly understood, especially in crop species under pathogen attack (Sanchez-Lucas and Luna, 2025).

Mechanisms underpinning immunity have been mainly characterised in isolation, whereas infections in natural environments frequently occur in combination with other stress factors simultaneously or sequentially, a threat aggravated by climate change. Intriguingly, multifactorial stress tends to impose divergent immune phenotypes to those evoked by each stressor when applied separately (Bidzinski *et al*., 2016; Suzuki *et al*., 2014; Atkinson *et al*., 2013; Prasch and Sonnewald, 2013; Son and Park, 2022; Zarattini *et al*., 2021). Given the rapid increase in atmospheric CO₂ may exceed the adaptive capacity of plants to fully reconfigure their regulatory and metabolic networks, it is of utmost interest to evaluate whether eCO2 conditions may potentially result in a suboptimal cue under future climate scenarios.

Importantly, the configuration of the immune states in plants have been achieved by rewiring in regulatory networks and the recruitment of alternative features according to selective pressures to co-ordinately counteract adverse cues efficiently (Fusco and Minelli, 2010; Dreze *et al*., 2011; He *et al*., 2020). Moreover, immune responses are articulated via singularities at different molecular layers, especially between transcriptomes and metabolomes (Garcia-Molina and Pastor, 2023). As result, attempts to manipulate plants for individual components to confer resilience to pathogens in disregard of the context in the network cause unexpected consequences (Liu *et al*., 2019; Ke *et al*., 2020). Similarly, strategies based on translating beneficial features identified in tractable plants into crop species tend to fail as the architectures of homeostatic networks are refined by singularities of each system and may drive to interdependences, the so-called interspecific barriers (Garcia-Molina and Leister, 2020; Kromdijk *et al*., 2016). In the same line, our previous works reported on how phosphate and Fe treatments provoke antagonistic immune phenotypes in Arabidopsis and rice (Garcia-Molina *et al*., 2025; Val-Torregrosa *et al*., 2022; Campos-Soriano *et al*., 2020). In consequence, systemic approaches deciphering networks underlying responses to each pathosystem in species of agronomical interest, and considering eCO2 conditions, are necessary to obtain a holistic picture about plant immunity in the years to come and to identify novel immune features to design strategies for resilience without risk of payoffs.

Tomato (*Solanum lycopersicum*) is a primary vegetal crop worldwide, widely cultivated for its nutritional and economic importance, that has been adopted as a model to study plant responses to eCO2. Increased CO2 availability tends to favour tolerance to abiotic stressors, principally heat and drought (Wang *et al*., 2024; Zhou *et al*., 2022; Jensen *et al*., 2024; Li *et al*., 2021). Conversely, its impact on interactions with pathogens depends on the pathosystem, as it has been reported to increase the susceptibility to the necrotrophic fungus *Botrytis cinerea*, the oomycete *Phytophthora parasitica* and the nematodes *Helicoverpa armigera*, *Meloidogyne incognita*, but reinforces defence against the bacteria *Pseudomonas syringae* and the tobacco and tomato mosaic viruses (Marino *et al*., 2025; Guo *et al*., 2012; Li *et al*., 2015; SUN *et al*., 2010; Jwa and Walling, 2001; Zhang *et al*., 2015; Hu *et al*., 2023; Li and Ahammed, 2023).

Tomato is frequently challenged by *B. cinerea*, a widespread pathogen responsible for grey mould disease that causes significant yield losses and negatively impacts on post-harvest quality (Petrasch *et al*., 2019; Blanco-Ulate *et al*., 2016). Defence against necrotrophic relies on complex metabolic and hormonal pathways, including jasmonate- and ethylene-mediated responses, which are tightly linked to the plant’s metabolic status and energy balance. Despite its agronomic relevance, few studies have addressed how tomato immunity will be affected under elevated CO₂ conditions. Here, we investigated how elevated CO₂, according to moderate projections for the end of the 21^st^ century (650 ppm), in combination with the associated temperature increase (+5 °C) (Intergovernmental Panel on Climate Change (IPCC), 2023; Meinshausen *et al*., 2020; Adak *et al*., 2023; Valone, 2021), influences defence responses in eight different tomato varieties during infection by *B. cinerea*. Our results revealed enhanced tolerance to *B. cinerea* across tomato accessions under eCO2. Systems-level analyses of metabolome and transcriptome during infection, together with the deconstruction of underlying gene regulatory networks (GRNs), revealed an uncoupling between transcription and metabolite accumulation. Furthermore, we isolated a central regulatory module sustained by WRKY and ERFs TFs articulating a transcriptional growth-defence trade-off. However, GRN perturbation through silencing sets of TFs in the central regulatory module highlighted the importance of basal mechanisms in counteracting *B. cinerea* infection, with outcomes largely dependent on genotype and CO2 availability. In this line, further metabolome analyses revealed that eCO2 led to reinforcement in structural and chemical barriers.

Thus, this work underscores the importance of considering multi-level network dynamics when interpreting plant stress responses and highlights the need for integrative approaches to understand how these responses are shaped under near-future environmental conditions.

## Results

### Elevated CO2 and temperature reinforce tomato tolerance to *Botrytis cinerea*

To assess the impact of climate change on tomato immunity against fungal infection, an experimental design was established to reflect moderate end-of-century projections of increased temperature (+5 °C) and atmospheric CO2 concentration (650 ppm) (Intergovernmental Panel on Climate Change (IPCC), 2023; Meinshausen *et al*., 2020; Adak *et al*., 2023). Tomato plants were grown either under environmental CO2 concentration (approx. 420 ppm) with a 16 h light/8 h dark cycle at 25 °C and 120 µmol photons m^−2^ · s^−1^ during the day and 18 °C at night (control), or under elevated CO2 (650 ppm) with corresponding day/night temperatures of 29 °C and 22 °C (eCO₂). Fungal bioassays were conducted with the necrotrophic pathogen *Botrytis cinerea*. 14-d-old tomato plantlets were systemically inoculated, and disease symptoms in aerial tissues were evaluated at 3 dpi. A diverse panel of tomato varieties was included in the study: Ailsa Craig (A), Better boy (BB), Coeur de boeuf (CB), Marmande (M), microTom (mT), Moneymaker (MM), Roma VF (R) and San Marzano (SM). Trypan blue staining of inoculated leaves revealed that *B. cinerea* infection caused substantial tissue damage under control conditions, although the severity varied across tomato varieties (**Figure 1A**, upper panel). Quantification of fungal biomass showed that BB, M and SM displayed high susceptibility. A and R exhibited moderate susceptibility, whereas CB, mT, and MM showed the lowest biomass levels and were classified as tolerant (**Figure 1B**). In contrast, eCO₂ markedly restricted fungal progression (**Figure 1A**, lower panel). Fungal biomass was significantly reduced by approximately three- to nine-fold compared to control conditions and converged towards the levels observed in the tolerant varieties (CB, mT, MM) (adjusted *P* ≤ 0.05, LSD’s test) (**Figure 1B**). Thus, despite intrinsic defences in basal resistance among tomato varieties, eCO2 consistently enhanced plant tolerance to *B. cinerea*.

**Figure 1.**
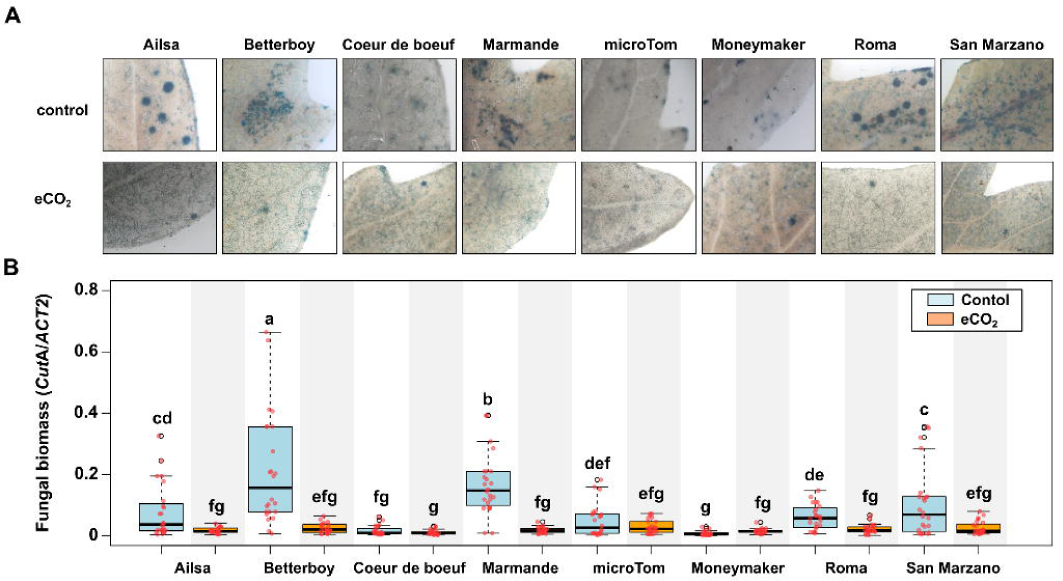
Immune phenotype of tomato varieties against *Botrytis cinerea*. 14-d-old tomato plantlets cultivated under control (16h light 25°C / 8h dark 18°C, 420 ppm CO2) and elevated CO2 environments (16h light 29°C / 8h dark 22°C, 650 ppm CO2, eCO2) were sprayed with suspensions of *Botrytis cinerea* spores (10^6^ spores/mL) and evaluated 3 dpi. (A) Leaf damage was monitored by trypan blue staining. Representative pictures are provided. (B) Fungal biomass was determined by qPCR with specific primers. Letters indicate statistically significant differences (LSD test, adjusted *P* ≤ 0.05, n = 10).

### Tomato varieties display heterogeneous metabolome reconfigurations in response to fungal infection and eCO2

To obtain a comprehensive view of molecular changes associated with *B. cinerea* infection under different growth conditions, we profiled the metabolomes of fungal infected and mock-treated plants grown under control and eCO2 across eight varieties using untargeted LC-MS/MS. This approach yielded a coverage of 9,102 compounds (3,127 in ESI- and 5,975 in ESI+) (**Table S1**). To capture the most prevalent condition-dependent responses, metabolome datasets were filtered for significantly accumulated metabolites (SAMs) associated with the interaction between the growth condition (control/eCO2) and treatment (mock/infection) (*P* ≤ 0.05, linear mixed models) (**Table S2**). Notably, the number of SAMs did not correlate with the immune phenotypes. For instance, the highest and lowest SAM counts were observed in BB (susceptible) and CB (tolerant) (873-936 SAMs), whereas intermediate numbers were detected in A, M and SM (300-400 SAMs), all of which were susceptible (**Figure 2A**). Furthermore, overlap of SAMs among varieties was minimal, indicating that each variety deploys distinct metabolome reconfigurations in response to fungal infection under eCO2 (**Figure S1A**).

**Figure 2.**
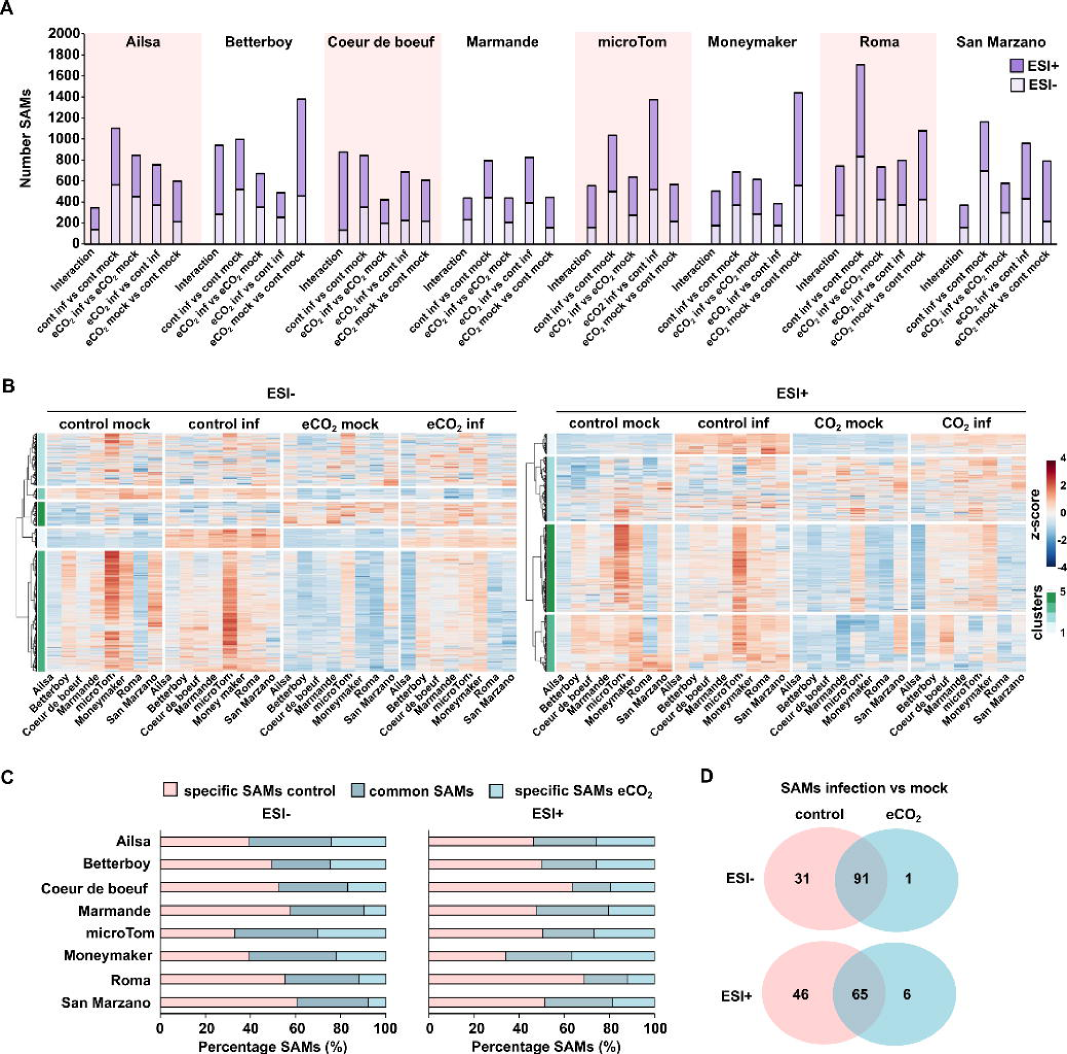
Metabolome response of tomato varieties against *Botrytis cinerea* under different growth conditions. 14-d-old tomato plantlets cultivated under control (16h light 25°C / 8h dark 18°C, 420 ppm CO2) and elevated CO2 environments (16h light 29°C / 8h dark 22°C, 650 ppm CO2, eCO2) were either mock treated or inoculated with suspensions of *Botrytis cinerea* spores (10^6^ spores/mL) and harvested 1 dpi. Metabolome profile was conducted by untargeted LC-MS/MS. (A) Bar plots indicate the number of significantly accumulated metabolites (SAMs) for the interaction between the condition (control/eCO2) and treatment (mock/infection) (*P* ≤ 0.05, linear mixed models, n=6) and pair-wise comparisons (adjusted *P* ≤ 0.05, Tukey’s test in linear mixed models, n=6) detected in ESI- and ESI+. (B) Heatmaps with hierarchical clustering according to the Ward D2 method for significantly changing compounds due to the interaction of factors in at least one variety. The median of log2-transformed normalised compound abundance was used to calculate z-scores and clusters were defined by the most optimal *k*-means partitioning. (C) Determination of fractions of common SAMs between growth conditions per variety. (D) Overlap between common SAMs responding to fungal infection in both growth conditions.

To visualise these divergences, the median abundance of metabolites identified as SAMs as result of the interaction in at least one variety were used to generate hierarchical clustering heatmaps for both ESI- and ESI+ compounds. As shown in **Figure 2B**, most compounds exhibited highly variable trajectories across varieties and tended to cluster primarily according to growth condition (control/eCO2), rather than genotype or treatment alone. Only cluster 1 (263 and 221 metabolites in ESI- and ESI+, respectively) displayed a consistent immune-responsive pattern, with stronger accumulation under control conditions compared to eCO2 (**Figure 2B**). The tentative annotation of compounds in cluster 1 is provided in **Table S3**. KEGG enrichment analysis revealed a significant overrepresentation (adjusted *P* ≤ 0.05, Fisher’s exact test) of pathways mainly related to primary and secondary metabolism, including carbohydrate and central carbon metabolism (e.g., pentose phosphate pathway, pyruvate metabolism, TCA cycle), amino acid biosynthesis, as well as pathways associated with secondary metabolites and hormone signalling. Notably, pathways linked to stress, such as ascorbate metabolism, plant hormone signal transduction, namely abscisic acid, and specialised defence compounds such as alkaloids were also enriched (**Table S4**).

To further dissect metabolome dynamics, additional SAMs were determined for biologically relevant pair-wise contrasts (adjusted *P* ≤ 0.05, Tukey’s test in linear mixed models), including: (i) infection vs mock under control conditions, (ii) infection vs mock under eCO₂, (iii) infection under eCO₂ vs infection under control, and (iv) mock under eCO₂ vs mock under control (**Table S2**). As observed for the interaction term, the number of SAMs varied widely across contrasts and varieties and did not correlate with susceptibility/tolerance to *B. cinerea* (**Figure 2A**). For instance, under control conditions 800-1,200 SAMs were detected in response to infection across most varieties, except for MM (tolerant) and R (susceptible) (600 and 1700 SAMs, respectively). Under eCO2 infection-responsive SAMs ranged from 660 to 800 in most varieties, except for CB (tolerant) and M (susceptible), which showed lower responses (around 400 SAMs) (**Figure 2A**). Consistent with the enhanced susceptibility observed under control conditions, 40-60% of infection-responsive SAMs were uniquely detected under control, whereas 20-35% were shared between control and eCO2 (**Figure 2A**). Strikingly, only a small core set of metabolites responded consistently to infection across all varieties (122 and 111 SAMs under control; 92 and 71 SAMs under eCO2 for ESI- and ESI+, respectively) (**Figure 2C** and **S1B**). Moreover, metabolite changes driven solely by growth conditions (mock eCO₂ vs mock control) or by infection across conditions (infection eCO₂ vs infection control) were comparable in magnitude to infection-induced responses and exhibited similarly high heterogeneity and low overlap among varieties (**Figure 2D and S1C-D**).

Collectively, these results reveal extensive genotype-dependent variability in metabolome composition and responsiveness, highlighting a strong context dependency of metabolic reprogramming. This high degree of heterogeneity precludes the straightforward identification of a conserved metabolic signature underlying immunity to *B. cinerea*, suggesting instead that multiple, genotype-specific metabolic strategies contribute to disease outcomes under different CO₂ environments. Nevertheless, an overall activation of a broad metabolic and regulatory response occurs upon *B. cinerea* infection, which is more attenuated under eCO2.

### Transcriptome profiling links gene expression reprogramming to varietal immune phenotypes

To gain deeper insight into the molecular responses underlying tomato plants immunity to *B. cinerea*, transcriptomes of M and SM (susceptible) and CB and MM (tolerant) were profiled. RNA-Seq datasets were used to identify differentially expressed genes (DEGs) with absolute log2(FC) ≥ 1 (adjusted *P* ≤ 0.05, Wald’s test) for both the interaction term and relevant pair-wise contrasts (**Table S5**). In contrast to metabolome profiles, susceptible varieties (M and SM) exhibited markedly higher number of infection-responsive DEGs compared to tolerant genotypes (CB and MM), particularly for the interaction term (approx. 1,000-1,600 vs 60-80 DEGs) (**Figure 3A**). Approx. 500 DEGs were shared between the two susceptible varieties, representing about half and one-third of the total interaction-responsive DEGs in M and SM, respectively (**Figure S2A**), suggesting a partially conserved transcriptional response associated with susceptibility.

**Figure 3.**
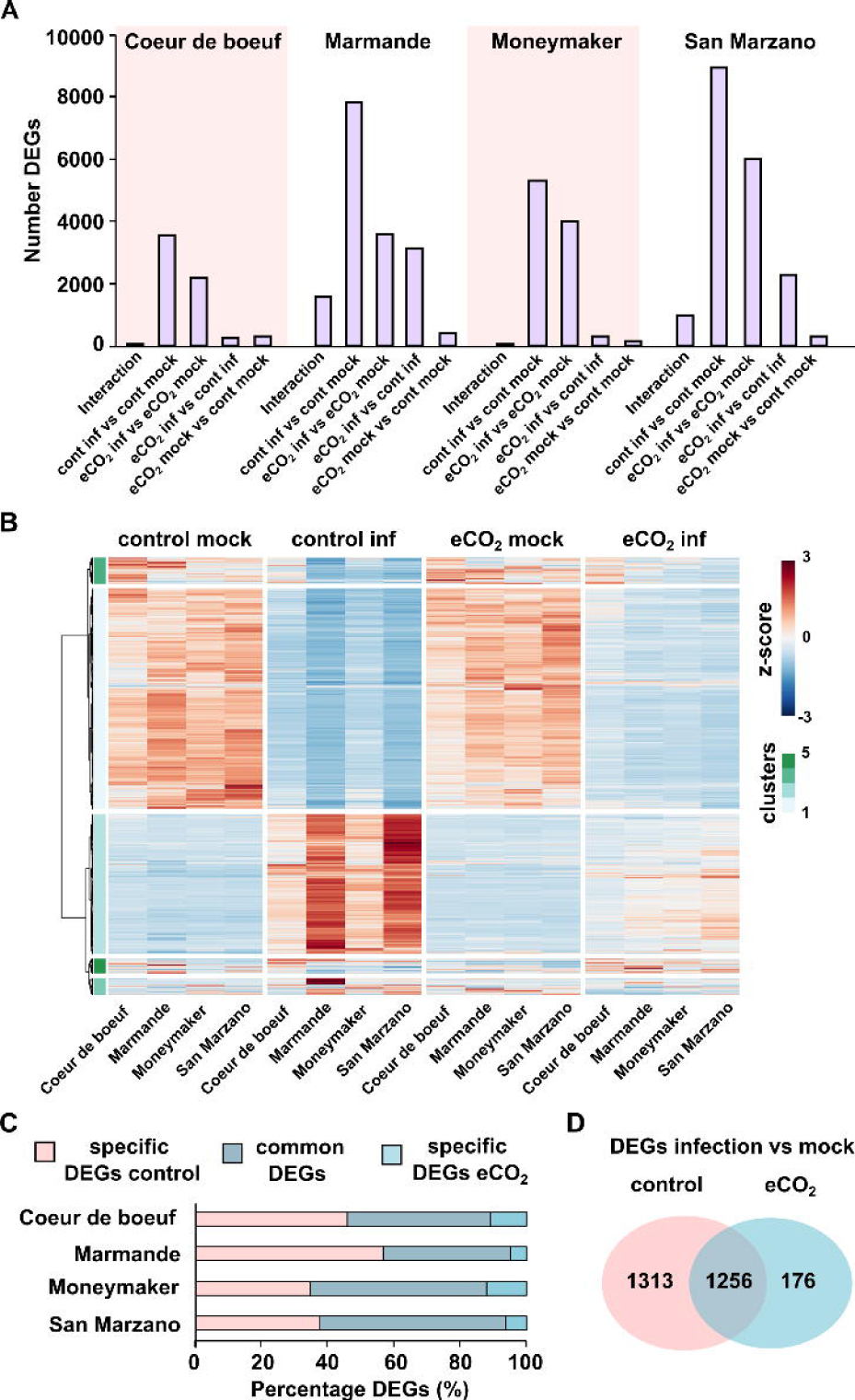
Transcriptome response of selected tomato varieties against *Botrytis cinerea* under different growth conditions. 14-d-old tomato plantlets cultivated as in Figure X were used for transcriptome profiling by RNA-Seq. (A) Bar plots indicate the number of differentially expressed genes (DEGs) for the interaction between the condition (control/eCO2) and treatment (mock/infection) and pair-wise comparisons (adjusted *P* ≤ 0.05, Wald’s test, n=3). (B) Heatmaps with hierarchical clustering according to the Ward D2 method for DEGs as result of the interaction of factors in at least one variety. The median of normalised counts was used to calculate z-scores and clusters were defined by the most optimal *k*-means partitioning. (C) Determination of fractions of common DEGs between growth conditions per variety. (D) Overlap between common DEGs responding to fungal infection in both growth conditions.

To identify dominant transcriptional patterns, normalised expression values for transcripts responding to the interaction term in at least one variety were used to generate a hierarchical clustering heatmap. These analyses did not reveal clear antagonistic expression programs between susceptible and tolerant varieties. Instead, steady-state levels of infection-responsive transcripts displayed amplified repression (cluster 1 and 4) and induction (cluster 2) in susceptible varieties (M and SM), particularly under control environment (**Figure 3B, Table S6**). Gene ontology (GO) enrichment analysis (adjusted *P* ≤ 0.05, Fisher’s exact test) of downregulated transcripts (1,023) revealed a marked repression of photosynthetic and chloroplast-associated processes, including chlorophyll biosynthesis, photosystem assembly and repair, the reductive pentose phosphate cycle, and photorespiration, together with broad inhibition of plastid organization and gene expression. Central carbon metabolism was also affected, as reflected by the downregulation of glycolysis, gluconeogenesis, and starch and carbohydrate metabolic processes (**Table S7**). In contrast, upregulated transcripts (579) were predominantly associated with stress and redox-related responses, including ROS detoxification, as well as amino acid and phenylpropanoid metabolism, indicating a shift from photosynthetic activity towards stress adaptation and secondary metabolism (**Table S7**). Notably, the magnitude of this transcriptional response was attenuated under eCO2 and resembled that of tolerant varieties (**Figure 3B**). These results suggest that repression of primary carbon metabolism and chloroplast functions is a defining feature of susceptibility, whereas maintaing these processes may contribute to counteract *B. cinerea* infection.

Consistent with the immune phenotypes, all varieties displayed 1.5 to 2 times more infection-responsive DEGs (infection vs mock) under control compared to eCO2 (approx. 3,500 vs 2,200 in CB, 7,800 vs 3,600 in M, 5,300 vs 4,000 in MM and 9,000 vs 6,000 in SM) (**Figure 3A**). Notably, susceptible varieties (SM and M) displayed tenfold higher number of condition-dependent DEGs compared to tolerant varieties (MM and SM) when comparing infection under eCO2 vs control (approx. 2,300 and 3,200 vs 160 and 300 DEGs) (**Figure 3A**), indicating a stronger transcriptional sensitivity to environmental conditions. Despite this variability, approx. 50% of infection-responsive DEGs were shared between control and eCO2 conditions within each variety, whereas eCO2-specific DEGs accounted for less than 10%. (**Figure 3C**), pointing to a conserved core response to fungal infection. Across varieties, 2,500 DEGs were commonly regulated under control conditions and 1,400 under eCO2, with a core-set of 1,200 DEGs shared between both environments (**Figure 3D**, **S2B and Table S8**). Functional annotation of this conserved set recapitulated the major biological processes identified for the interaction of factors, reinforcing their central role in the immune response (**Table S9**). In contrast, the direct impact of eCO2 on the transcriptome under mock conditions was modest, with only 160-400 DEGs detected across varieties (**Figure 3A**). Collectively, these findings indicate that tomato plants mount a conserved core transcriptional defence response against *B. cinerea*, the magnitude of which scales with susceptibility, while additional layers of transcriptional reprogramming are condition-dependent and modulated by eCO2 availability.

### Integrated transcriptomic and metabolomic analysis reveals CO₂-dependent decoupling and attenuation of genotype-specific responses to *B. cinerea* infection

Our analysis of transcriptomic and metabolomic datasets revealed a marked discordance between transcriptional regulation and metabolite accumulation across interconnected metabolic pathways. Transcriptomic data showed strong repression of photosynthetic and chloroplast-related processes, including photosystem assembly, chlorophyll biosynthesis, the Calvin cycle, and photorespiration, alongside downregulation of central carbon metabolism (glycolysis/gluconeogenesis, sucrose biosynthesis and glycine decarboxylation), cellular redox homeostasis and growth-associated functions (**Figure 4A**). Strikingly, this transcriptional repression was accompanied by the accumulation of intermediates in key central carbon pathways such as the TCA cycle, pyruvate metabolism, and the pentose phosphate pathway, as well as increased levels of amino acids, secondary metabolites, and antioxidant compounds such as ascorbate (**Figure 4A**). This apparent decoordination suggests that, despite broad transcriptional downregulation of primary metabolism aimed at restraining growth and biosynthetic capacity, metabolic fluxes are maintained or redirected towards a survival-oriented state, consistent with stress activation. These patterns point to the emergence of system-wide metabolic bottlenecks rather than true pathway activation, whereby reduced transcriptional support limits enzymatic capacity, constrains metabolic flux and, ultimately, leads to the accumulation of intermediates. Nevertheless, transcriptome and metabolome changes converged on increased stress-related features (**Figure 4A**).

**Figure 4.**
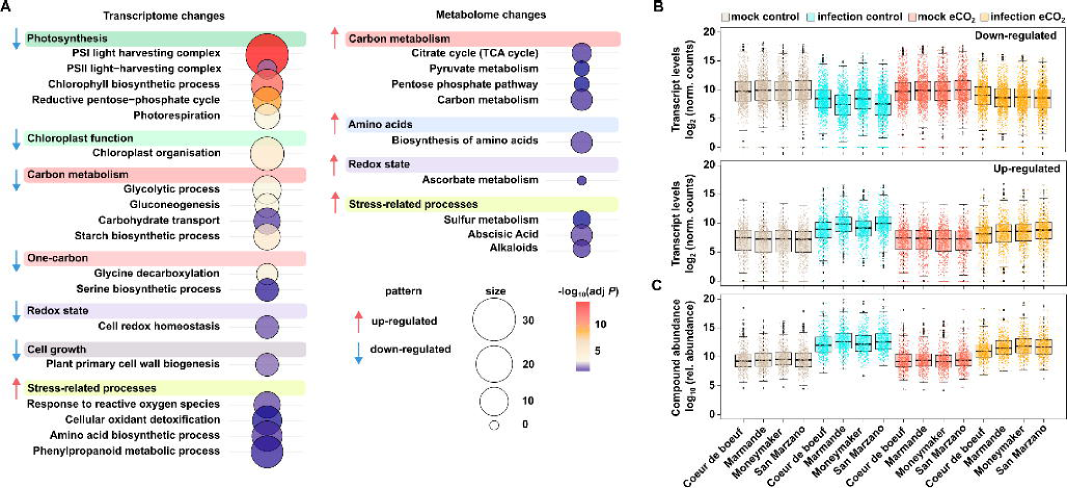
Integrated transcriptomic and metabolomic model of CO2-dependent responses to B. cinerea. A) Summary of Gene Ontology term enrichment for transcriptomic changes and KEGG pathway enrichment for altered metabolites (adjusted *P* ≤ 0.05, Fisher’s exact test). Terms are grouped by category. Red and blue arrows indicate up- and down-regulation of the associated features, respectively. Circle size and colour are proportional to the number of hits and the -log10-transformed adjusted P, respectively. Bar plots depict the distribution of steady-state levels of down- regulated transcripts (cluster 2) and up-regulated transcripts (clusters 1 and 4) (**B**), as well as metabolite relative abundance (cluster 1) (**C**), corresponding to the heatmaps shown in Figures 2 and **3**.

Notably, under control conditions, the transcriptional responses displayed strong genotype-dependent variability. Susceptible varieties exhibited a markedly amplified transcriptional repression compared to tolerant genotypes, consistent with a more severe disruption of metabolic coordination. In contrast, tolerant varieties maintained a more moderated response, suggesting a greater capacity to preserve metabolic balance under infection (**Figure 4B, C**). By comparison, eCO₂ importantly attenuated both transcriptional repression and metabolite accumulation across pathways. Under these conditions, responses became more homogeneous across genotypes, with susceptible varieties converging towards the attenuated profiles observed in tolerant plants (**Figure 4B, C**). These observations indicate that eCO₂ not only reduces the magnitude of infection-induced reprogramming but also dampens genotype-dependent variability, suggesting that increased carbon availability buffers the system against the emergence of divergent metabolic states.

Together, these findings support a model in which CO₂ availability governs both the magnitude and variability of plant responses to infection, determining whether transcriptional reprogramming leads to metabolic dysfunction or to to a buffered, functionally integrated defence response.

### Progressive assembly of Gene Regulatory Networks reveals a conserved backbone with condition-specific metabolic layering

To further understand how transcriptional regulation underpins the metabolic reprogramming observed during *B. cinerea* infection, gene regulatory networks (GRNs) were constructed based on core basal defence genes found under control and eCO2 conditions across all varieties (basal GRN, GRNb), only under control condition (GRN under control, GRNc) or those prevalent in susceptible varieties (interaction term, susceptible GRN, GRNs) (**Figure 3D**). GRNs were inferred from RNA-Seq datasets using the *GENIE3* algorithm and network density ajusted with *DIANE* (Huynh-Thu *et al*., 2010; Cassan *et al*., 2021) (see Methods section).

Topologically, the three resulting GRNs (**Figure 5A** and **Table S10**) revealed a progressive expansion in scale and size characterised by the sequential incorporation of TFs (from 62 in GRNb to 116 in GRNc) and target nodes (from 1,008 in GRNb to 1,829 in GRNc) (**Figure 5B**). Normalised TF connectivity in GRNb ditributed as in natural-systems networks, i.e., being few TFs highly connected while the majority displayed lower degree (Barabási and Albert, 1999) (average degree 0.3) with most nodes targeted by four TFs (**Figure 5C, D**). To note, newly incorporated TFs in GRNc and GRNs displayed predominantly low-to-moderate connectivity, as the average connectivity decreased to 0.2 and 0.15, respectively, and the targets connected to two to three TFs (**Figure 5C, D**). Furthermore, almost half of the TFs already present in GRNb established connections with novel recruited nodes for GRNc and vice versa. Conversely, around 45% of the incorporated TFs in the GRNc established interactions with pre-existing nodes in GRNb. A similar pattern was found in the case of the GRNs (**Figure 5E**). Thus, our analysis indicate that GRNs architectures remained highly structured resulting in a conserved backbone topology that anchors the network and points to potential constraints preventing excessive expansion. This suggests that infection-responsive regulatory programs are not rewired *de novo* but rather built upon a stable core of basal defence components. The progressive recruitment of TFs with limited connectivity suggests a distributed regulatory strategy in which multiple weakly connected regulators fine-tune existing network modules rather than establishing dominant hubs.

**Figure 5.**
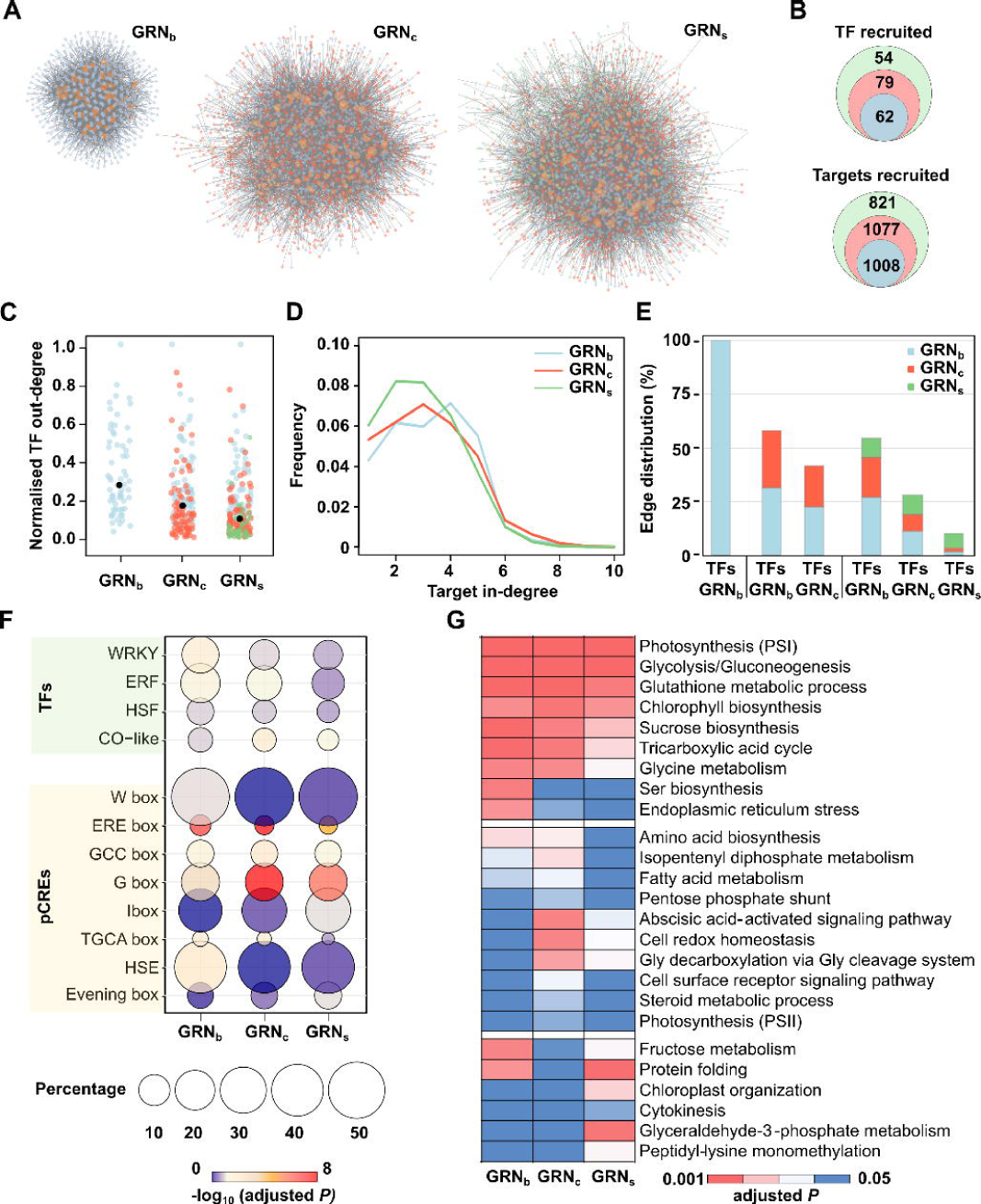
Gene Regulatory Network analyses. **A**) Networks underpinning basal immunity (GRNb), common defence mechanisms in control conditions (GRNc) and in susceptible varieties under control conditions (GRNs) were constructed using normalised RNA-Seq counts with the *GENIE3* algorithm, and network density was filtered using *DIANE*. **B**) Increase size and scale increase of the GRNs. **C**) Dot plots showing the connectivity of transcription factors (TFs) in each GRN. Outdegrees were normalised from 0 to 1, and average connectivity indicated by black dots. **D**) Density plots summarising the distribution in the number of nodes targeted per TFs. **E**) Bar plot showing the interconnectivity among layers in the GRNs. Bar fractions represent the connections between TF and targets, according to the network in which they were incorporated. **F**) Significant enrichment in TFs families (adjusted *P* ≤ 0.05, Fisher’s exact test) and putatitve cis-regulatory elements (pCREs) within 1,000 bp upstream promoter regions on the nodes (adjusted *P* ≤ 0.05, Z-score test) in each GRN. **G**) Functional annotation of nodes according to Gene Ontology biological processes terms (adjusted *P* ≤ 0.05, Fisher’s exact test) for each GRN.

Then, a focused analysis of TFs was performed to identify the key regulators defining the structure and dynamics of the GRNs. In GRNb, significantly enriched TFs (adjusted *P* ≤ 0.05, Fisher’s exact test) predominantly belonged to the WRKY, ERF, and HSF families, in addition to CONSTANS (CO)-like, whereas in GRNc the enrichment remains mainly associated with ERFs. The relevance of these TF families was diluted in the GRNs (**Figure 5F**). This distribution was consistent with the observed enrichment of putative cis-regulatory elements (pCREs) motifs in the promoters of the target genes (adjusted *P* ≤ 0.05, Z-score test) (**Figure 5F, Table S11**). Notably, the role of WRKY, ERF, and HSF families, as well-known modulators of stress responses, and the coherence between TF enrichment and promoter motif composition provides functional support for the inferred GRNs.

Functional annotation of successive network layers revealed a clear trajectory in biological processes. Basal structural network modules, those in GRNb , were enriched in processes related to photosynthesis and central carbon metabolism, including sucrose biosynthesis and the TCA cycle, as well as redox functions (**Figure 5G, Table S12**). As networks expanded, additional layers incorporated nodes associated with amino acid and lipid metabolism and stress hormones (ABA) in the GRNc, followed by further enrichment of pathways linked to stress responses and defence signalling (protein folding, chloroplast organisation) in the GRNs (**Figure 5G, Table S12**). This progression indicates a hierarchical reorganization of transcriptional control, in which primary metabolic functions are progressively redirected towards stress adaptation.

Together, these results support a model in which tomato transcriptional responses to infection are governed by a core regulatory backbone that is progressively modified through the addition of peripheral regulatory layers. This modular expansion enables the coordinated transition from primary metabolism to defence-associated processes, providing a mechanistic framework linking transcriptional regulation to the metabolic reprogramming observed under different genotypes and CO₂ regimes.

### Gene Regulatory Networks encompass a module that mediates a trade-off in resource allocation between growth and defence processes

The architecture of GRNs, characterised by a controlled recruitment of TFs, evoked a system that allows flexible adaptation to counteract pathogen infection by progressive implementation of biological processes depending on the severity of the threat on the plant. Then, we next aimed at delineating the key modulators within each GRN. Thereto, the fraction of putative *bona fide* targets per TF, based on the number of connected nodes harbouring their cognate pCREs, was used as a proxy for regulatory relevance. This approach identified three WRKY (Solyc02g072190, Solyc02g093050, Solyc02g032950) and three ERFs (Solyc09g089910, Solyc01g090340, Solyc08g078190; hereafter ERF1) consistently present across the three GRNs, as well as an additional pair of ERFs (Solyc04g078640, Solyc03g005520; hereafter ERF2) specifically for the GRNc and GRNs (**Table S13**). These TFs were used as seeds to extract their vicinity (**Table S14**), what rendered a subset of nodes organised in a module (**Figure 6A**). The module expanded both structurally and in complexity, increasing from 189 nodes and 194 edges in GRNb to 302 and 318 edges in GRNs (**Figure 6A**). Furthermore, the selected TFs exhibited enhanced interconnectivity, with interplays among members of the same family and between different families rising from 11 and 5 in GRNb to 24 and 15, respectively, in GRNs (**Figure 6A**). Collectively, these observations support a potential role of this regulatory module in coordinating plant defence.

**Figure 6.**
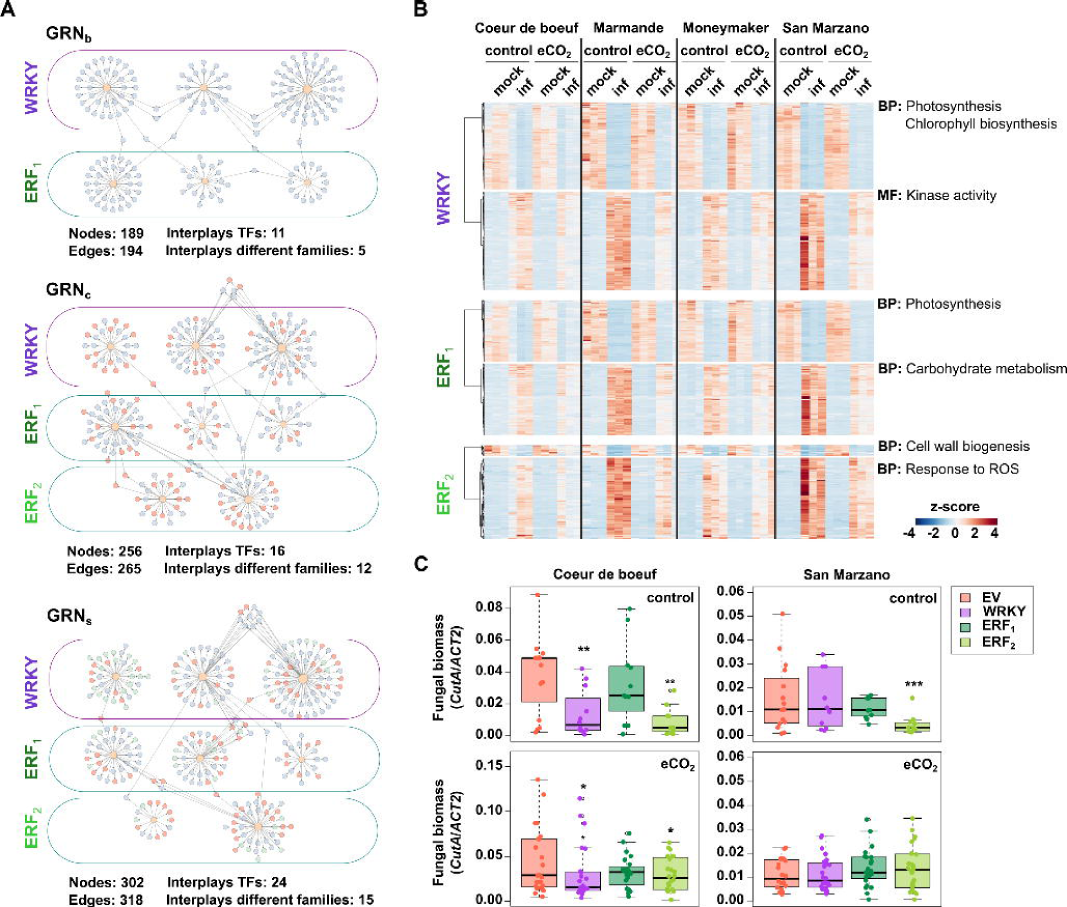
Participation of the central regulatory module in Gene Regulatory Networks in maintaining the trade-off between growth and defence. A) Cartoon depicting the subnetwork isolated with most relevant TFs in each GRN (WRKY, ERF1, ERF2). The topological metrics are provided. B) Heatmap with hierarchical clustering summarising the patterns of transcript levels of the targets of each group of TFs according to the WardD2 method. C) *B. cinerea* fungal biomass in Coeur de boeuf and San Marzano accessions at 3 dpi in plants cultivated under standard (control) or eCO2 conditions. VIGS were used to produce a control (empty vector, EV) or target TF groups (WRKY, ERF1, ERF2). Asterisks indicate significant differences to EV (* adjusted *P* ≤ 0.05; * adjusted *P* ≤ 0.01; *** adjusted *P* ≤ 0.001, Linear mixed models).

Interestingly, the steady-state levels of the WRKY and ERF1 targets displayed contrasting yet balanced patterns in response to the infection under both conditions (control and eCO2), although the intensity depended on the immune phenotype of the variety (**Figure 6B**). Functional annotation of transcripts that decreased in response to *B. cinerea* revealed enrichment in GO terms related to the biological process “photosynthesis” (WRKY and ERF1), “chlorophyl biosynthesis” (WRKY) and “cell wall biogenesis” (ERF2). Conversely, WRKY targets that increased upon infection were enriched in the molecular function term “kinase activity”, with many of these genes playing well-defined roles in immune responses (**Table S15**). ERF1 and ERF2 targets were associated with stress-related biological processes, including, “carbohydrate metabolism” and “response to ROS” (**Table S15**). Together, these findings indicate that this WRKY–ERF regulatory module co-ordinately activates defence programs while repressing primary metabolic pathways, thereby establishing a transcriptional growth-defence trade-off. Notably, this configuration appears to operate as a compensatory regulatory circuit that balances resource allocation rather than maximizing immune output. We therefore propose that additive regulatory features modulating this balance are particularly relevant in susceptible immune states, where this trade-off may constrain effective defence responses.

To assess the contribution of this regulatory module to plant immunity, a Virus-Induced Gene Silencing (VIGS) approach was employed to independently knock down distinct sets of TFs separately in the CB and SM cultivars (see Methods), representing tolerant and susceptible responses to *B. cinerea*, respectively. Silencing two out of the three WRKY TFs resulted in significantly reduced fungal biomass in CB plants under both control and eCO2 compared to the control group (empty vector) (**Figure 6C**). In contrast, VIGS targeting ERF2 TFs led to a marked reduction in fungal biomass in both CB and SM, although this effect was only observed under the control environment (**Figure 6C**). Silencing of ERF1 TFs did not affect fungal infection (**Figure 6C**). Taken together, these results suggest that WRKY and ERF2 TFs define a regulatory module that maintains a compensatory growth-defence balance, resulting in a partially constrained and suboptimal immune output. Perturbation of this module appears to relieve this constraint and enhance resistance, highlighting it as a potential target for improving plant immunity.

### Elevated CO2 promotes a metabolically primed state characterised by enhanced structural and chemical defences against *B. cinerea* in tomato

The emergence under eCO2 of a minimal yet efficient growth-defence trade-off, together with the enhanced tolerance displayed by CB and SM upon targeting central TFs within the identified regulatory module raised the question of whether reallocating resources towards constitutive defensive barriers could efficiently restrict *B. cinerea* progression. Because our metabolome annotation relied on internal libraries enriched in defence-related secondary metabolites (Gamir *et al*., 2014), the metabolomic datasets were re-examined focusing on defence-associated compounds. The infection-induced log2(FC) under both control and eCO2 conditions was calculated across the eight varieties (**Table S16**).

Infection-induced indole-derived metabolites, particularly the tryptophan-derived compounds IAA-Ile and 5- hydroxytryptophan, were consistently attenuated under eCO₂ (**Figure 7** and **S3**). Likewise, seveal metabolites associated with nucleotide turn-over, energy metabolism and redox metabolism, including adenosine, ADP, dAMP, NMN, NADP, and FMN, showed weaker accumulation trend during infection (**Figure 7** and **S4**). These changes suggested reduction in signalling activity and decreased reliance on energetically demanding regulatory responses under eCO₂. Consistently, eCO₂ also dampened the abundance of the antioxidant ergothioneine and the oxylipins hydroxy- and trihydroxy-octadecenoic acid (HODE and TriHOME), pointing to reduced activation of oxidative and lipid-derived defence pathways during pathogen challenge (**Figure 7** and **S5**).

**Figure 7.**
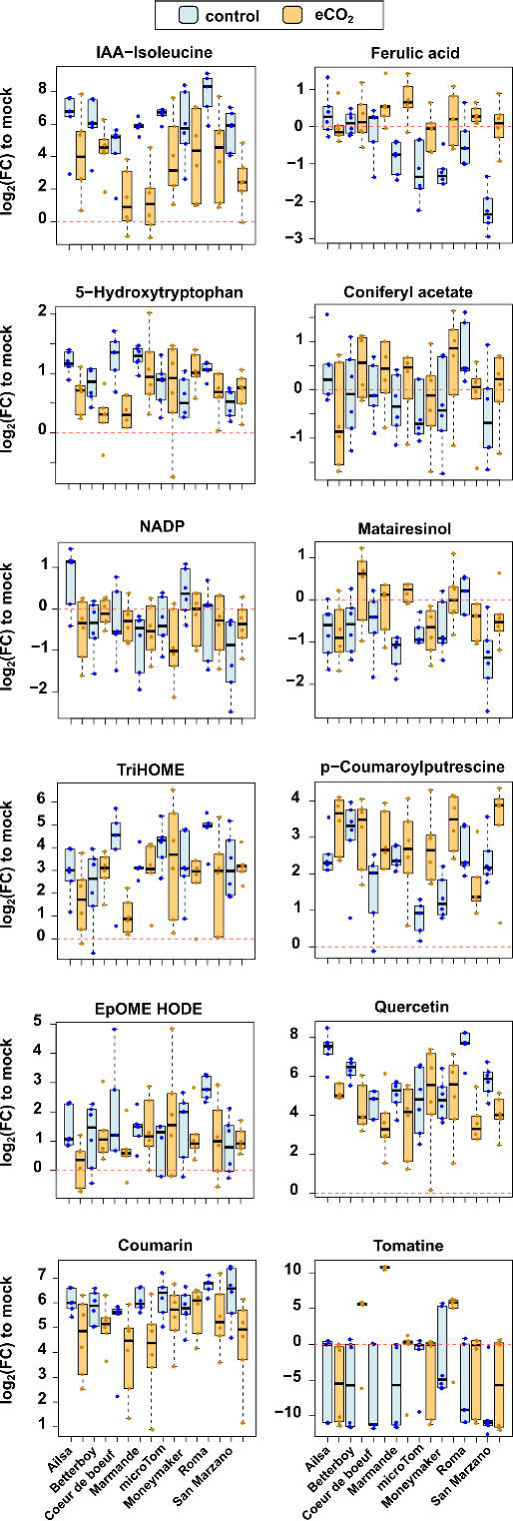
Representative metabolites illustrating the defence reprogramming induced by elevated CO₂ during *Botrytis cinerea* infection. Metabolome datasets were filtered for representative defence-related compounds. Boxplots depict log2-transformed fold-changes (FC) of infection-induced changes to mocks under control (light blue) and eCO₂ (light orange).

On the other hand, infection-induced accumulation of coumarin-related compounds (coumarin, umbelliferone, and glucosyl-coumarate) was reduced under eCO₂, whereas phenylpropanoids associated with cell-wall reinforcement and lignification, such as ferulic acid and coniferyl acetate, accumulated more strongly. This shift was further supported by the enhanced accumulation of the lignans matairesinol, pinoresinol glucoside and sesamolinol glucoside (**Figure 7** and **S6**), indicating a marked reallocation of carbon flux towards structural defence barriers upon infection under eCO₂. Additional evidence for this metabolic redirection was provided by the accumulation of p-coumaroyl- putrescine, a defence-associated phenolamide (**Figure 7** and **S6**), linking phenylpropanoid and polyamine metabolism, consistent with a greater investment in defence reinforcement and protective barriers. Regarding flavonoid metabolism, the anthocyanin pelargonidin-3-glucoside accumulated more strongly under eCO₂, while the induction of quercetin, hesperetin and isoquercitrin was reduced (**Figure 7** and **S7**). This pattern indicates a redistribution of metabolic flux within the flavonoid pathway rather than a global enhancement of flavonoid biosynthesis (**Figure 7** and **S8**). Finally, eCO₂ favoured the accumulation of specialised defence metabolites including, in addition to p-coumaroyl-putrescine, resveratrol, stearic acid and, particularly, tomatine, a major antimicrobial glycoalkaloid in tomato, strongly associated with pathogen resistance. Increased accumulation of these compounds further supports enhanced investment in chemical defence under eCO2 conditions (**Figure 7** and **S5**).

Collectively, our targeted metabolome analysis indicates that eCO2 reconfigures the metabolic landscape of tomato during *B. cinerea* infection, shifting defence responses from signalling, redox- and oxylipin-associated processes towards the accumulation of structural and chemical defensive compounds. Such a metabolic configuration would be indicative of a metabolically primed state, whereby tolerance would be achieved not through stronger inducible responses but through a greater reliance on phenylpropanoid-derived barriers and antimicrobial metabolites.

## Discussion

How tomato plants will cope with *B. cinerea* infection under environments associated with climate change remains insufficiently explored. In this study, we showed that eight tomato varieties grown under conditions simulating near-future scenarios (650 ppm CO2), combined with the consequent temperature increase (+5 °C), exhibited enhanced tolerance to *B. cinerea*, regardless of their baseline susceptibility (**Figure 1**). Our observations contrast with the findings reported by (Zhang *et al*., 2015). However, in their study elevated CO2 was applied only during the four days preceding the infection, and without an accompanying increase in temperature, thereby excluding the effects of long-term high CO2 exposure during plant development.

A primary consequence of sustained eCO2 is the increased carbon fixation through photosynthesis, ultimately leading to improved plant energetic states (Thompson *et al*., 2017; Kaiser *et al*., 2017). This is particularly relevant because the activation of plants defence programs is energetically costly and must be balanced against growth and reproductive demands (Wang and Wang, 2014; Zrimec *et al*., 2025). This trade-off is especially critic in crop species, as domestication has imposed strong genetic bottlenecks that have reshaped plant genomes to favour yield at expenses of adaptive potential. Indeed, pan-genome analyses have shown that domesticated species often display extensive presence–absence variation in resistance gene and lack fractions of the accessory genome present in their wild relatives, including genes associated with tolerance to biotic stress (Bourne *et al*., 2025; Rose *et al*., 2025; Ji *et al*., 2024; Amas *et al*., 2023). Accordingly, we propose that continuous exposure to eCO2 may provide an excess of energetic resources sufficient to alleviate trade-offs, thereby enabling a more efficient activation of defence responses against *B. cinerea*.

The metabolome profile of tomato plants infected with *B. cinerea* revealed high heterogeneity across genotypes and failed to discriminate susceptible from tolerant varieties (**Figure 2**). This lack of convergent patterns would be consistent with varietal differences in morphological traits, such plant height, leaf size or thickness and developmental stage. Nevertheless, we identified a discrete cluster of approximately 500 compounds (264 in ESI- and 222 in ESI+) that consistently increased due to the interaction between conditions (control/eCO2) and treatments (mock/infected) (**Figure 2B**). Tentative annotation of those metabolites pointed to the accumulation of intermediates from primary metabolism (carbon and amino acid pathways), as well as in redox-related and stress-associated compounds during infection under both conditions (**Figures 2**, **4**). However, the transcriptome landscape following *B. cinerea* infection revealed, besides induction of canonical stress responses such as secondary metabolism or oxidative stress alleviation, severe repression of chloroplast-associated processes, central carbon metabolism, glycine decarboxylation and cell growth (**Figures 3**, **4**). Notably, the intensity of transcriptome changes was greater under control conditions than under eCO2, particularly in M and SM (susceptible varieties) compared to MM and CB (tolerant varieties) (**Figure 4B**). Accordingly, although infection imposes a marked transcriptional shutdown of photosynthetic carbon assimilation, it simultaneously leads to accumulation of intermediates from the pentose phosphate pathway, the TCA cycle and amino acid metabolism, revealing a pronounced decoupling between gene expression and metabolic output (**Figure 4A**). Such accumulation of compounds likely reflects metabolic bottlenecks arising from reduced enzymatic capacity and/or impaired flux distribution, rather than pathway activation. In particular, the build-up of Gly and Ser, together with reduced glycine decarboxylation, suggests disruption of the photorespiratory–one-carbon interface, reinforcing the notion of flux restriction at key metabolic nodes. In contrast, eCO₂ attenuates these effects by sustaining photosynthetic input, mitigating carbon limitation, partially preserving the metabolic flux. These findings support a model in which the infection under control conditions drives the system into a metabolically constrained state characterised by reduced flux and accumulation of intermediates, depending on the innate immunity of the variety, while eCO2 governs the system-level capacity of the plant to maintain homeostasis while activating defence.

An important observation derived from our multi-omics analysis is that infection imposed a broadly similar transcriptome landscape across all genotypes and conditions, with the intensity of the reconfiguration emerging as the main distinguishing feature. In other words, different immune phenotypes do not appear to rely on entirely distinct sets of transcriptional changes but rather reflect differences in the functionality and/or efficiency of shared immune components depending on the genotype or condition. Alternatively, these phenotypic differences may involve the implementation of additional regulatory layers, such as the metabolome or basal immune features. From a systems perspective, GRN deconstruction revealed that transcriptional responses underlying the transition from resistant to susceptible defence states are orchestrated by a core regulatory backbone that is progressively modified through the addition of peripheral regulatory layers (**Figure 5**). This modular expansion of GRNs suggests that TF recruitment is involved in fine-tuning the magnitud of the immune response as the infetion progresses. Such dynamic regulation would enable a coordinated shift from repression of primary metabolism to activation of defence-associated processes, providing a mechanistic framework linking transcriptional regulation to the metabolic reprogramming. Notably, we identified a WRKY–ERF regulatory module that appears to coordinately repress primary metabolic pathways while promoting defence-related gene expression, thereby establishing a transcriptional switch contributing to the growth– defence trade-off (**Figure 6A,B**). However, silencing either WRKY or ERF2 TFs sets within this regulatory module enhanced tolerance to *B. cinerea* in CB (tolerant) under both CO2 conditions and in SM (susceptible) only under control conditions (**Figure 6C**). These results highlighted the existence of alternative innate defence strategies among tomato varieties that may be more efficient and/or energetically favorauble by attenuating the growth–defence trade-off. Consistent with this interpretation, the innerently stonger basal defence capacity of CB might explain why WRKY silencing enhanced the basal immunity under both conditions, while in SM it would not confer additional benefits under eCO2.

Our targeted metabolic analysis supported the hypothesis that tolerance acquired under eCO₂ is achieved through defence reprogramming rather than enhanced activation of inducible immune response. Specifically, eCO2 appears to redirect defence metabolism from canonical defence-signalling pathways towards the accumulation of structural and chemical defence components, favouring a metabolically primed state (**Figure 7**). Modification of cell wall composition and lignification through stimulation of phenylpropanoid metabolism under carbon availability is consistent with previous evidence in the literature to ultimately reinforce physical barriers against *B. cinerea* invasion (Singh, 2025; Miedes *et al*., 2014; Molina *et al*., 2024; Khan *et al*., 2026). Beyond lignification and phenylpropanoid-derived reinforcement, other structural barriers such as the cuticle and trichomes may also be reinforced under eCO₂. The cuticle acts as the first physical interface limiting *B. cinerea* entry, while trichomes and associated secretory compounds provide both mechanical and chemical protection to the fungus (Arya *et al*., 2021; Kannangara *et al*., 2007; Gutensohn *et al*., 2014; Zabel *et al*., 2021). In parallel, increased carbon availability is reported to promote the accumulation of secondary metabolites, including phytoalexins, alkaloides and terpenoids, which exhibit antimicrobial activity and further restrict *B. cinerea* infection (Devadze *et al*., 2025; Qaderi *et al*., 2023). Consequently, these observations support the working model we propose in **Figure 8**.

**Figure 8.**
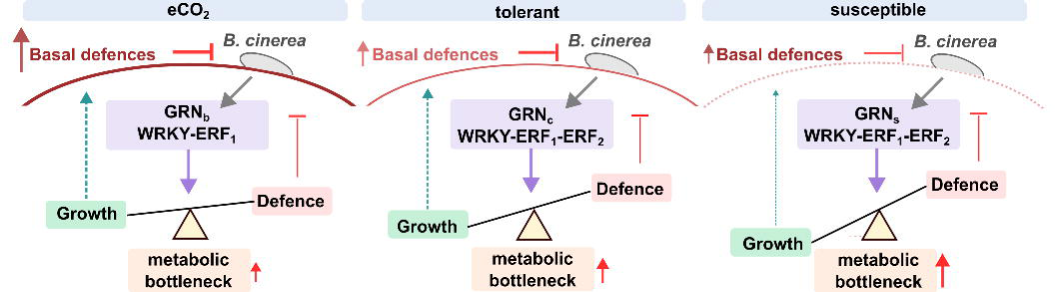
Model for the interplay between growth conditions and tomato immune responses to *B. cinerea*. Tomato plants grown under elevated CO2 conditions (eCO2; 650 ppm CO2 and +5 °C) exhibit enhanced carbon fixation, enabling an optimal metabolic state and the establishment of efficient basal defences. Infection by *B. cinerea* would activate a basal configuration of the Gene Regulatory Network (GRNb) to adjust the growth-defence trade-off to minimise metabolic bottlenecks to mount additional immune layers, i.e., structural and chemical barriers. This state would not interfere with allocation of energy to basal defence. In tolerant varieties, basal barriers are present, but infection would trigger a higher-order GRN (GRNc) that partially arrests primary metabolism while inducing complementary defence responses. Despite an increased accumulation of intermediate metabolites, this scenario would not substantially compromise sustaining basal defence. In contrast, susceptible varieties possess weak basal defence, and infection would activate a superior level in the GRN (GRNs) to drive adaptive responses. This would lead to severe growth repression and a strong arrest of metabolic fluxes, ultimately preventing the effective reinforcement of basal defences and the necessity to deploy an inducible signalling pathway, such as indole- and oxylipin-mediated responses.

As discussed above, cultivated tomato varieties have likely lost part of their defensive genetic repertoire during domestication. In this context, focusing on structural components or fundamental regulatory/signalling features may provide a more robust strategy to enhance resistance across diverse genetic backgrounds. Then, determining the relative contribution of pre-existing structural and chemical barriers to *B. cinerea* restriction under conditions of energetic surplus will be an imporant avenue for future investigation.

Overall, our study highlights how modern crops harbour finely tuned regulatory modules that sustain critical growth-defence trade-offs throuhg complex interplays across multiple molecular layers. Dissecting these mechanisms and understanding how they are coordinated across regulatory, transcriptional and metabolic levels will be essential to guide breeding strategies aimed at improving plant immunity, particularly under future climate change conditions.

## Methods

### Plant growth cultivation

*Solanum lycopersicum* seeds for Ailsa Craig, Better boy, Coeur de boeuf, Marmande, Moneymaker, microTom, Roma VF and San Marzano varieties propagated at CRAG or UJI were used in this study. For experiments with plantlets, seeds were sown on Jiffy-7 substrate (Jiffy Group International, Zwijndrecht, the Netherlands) under a 18 h light (25°C and 120 μmol m^−2^ s^−1^)/8 h dark (18°C) cycle with a relative humidity of 65% and atmospheric CO2 concentration (approx. 420 ppm) in FitoClima 600 PLH LED cabinets (Aralab, Rio de Mouro, Portugal). Treatments for eCO2 consisted of raising CO2 concentration to 650 ppm and increasing temperature cycles to 29°C/22°C. For experiments with adult plants, sees were cultivated in pots with a mixture of peat and vermiculite (3:1) for four to five weeks under the same condition as described above. Watering was conducted three times per week.

### Fungal infection assays

*Botrytis cinerea* was grown on full potato dextrose agar medium at 25°C in darkness for 15 days. Fungal spores were prepared and diluted at 1 ×·10^6^ spores ml^−1^ on ¼ potato dextrose broth. Spore suspensions were surface applied. Cell damage was visualised by trypan blue staining as in (Salguero-Linares *et al*., 2022). Fungal biomass was quantified from entire leaves at 3 dpi. Approximately 10 ng of gDNA extracted with Edward’s buffer were prepared for real-time qPCR with SYBR Green I dye using specific primers for *B. cinerea CutA* and the tomato *ACTIN2* (*ACT2*) (**Table S17**) in a Roche Light Cycler 480 instrument (Roche, Basel, Switzerland). The two-step protocol consisted of an initial cycle at 95°C for 10 min and 45 cycles of 95°C for 10 s and 60°C for 20 s. Fungal biomass was determined according to the 2^−Δ(Ct^ *^CutA^*^− Ct^ *^ACT2^*^)^ method. Outliers were detected according to the Interquartile Range method (2.5 x IQR) and replaced by imputation according to the *k*-nearest neighbours (*k* = 5) with the *VIM package* in RStudio. Experiments were conducted in biologically independent triplicates with n > 8 plants. Statistical differences were calculated using linear mixed models.

### Transcriptome and metabolome profiling and analysis

Samples for high-throughput profiling were prepared from rosette leaves harvested 8 h after the onset of light and flash frozen in liquid nitrogen. RNA-Seq and untargeted LC-MS/MS metabolomics were conducted as reported in Garcia-Molina and Pastor, 2024 with adaptations for tomato. Briefly, total RNAs (n = 3 independent biological replicates) were isolated with the Maxwell RSC Plant RNA Kit (Promega, USA). RNA-seq libraries were prepared and paired-end sequenced (2 × 150 bp) by Novogene (Munich, Germany) with standard Illumina protocols and analysed with Trimmomatic/RNA STAR/featureCounts/DESeq2 pipeline (Bolger *et al*., 2014; Dobin *et al*., 2013; Liao *et al*., 2014; Love *et al*., 2014) using the *Solanum lycopersicum* ITAG4.0 reference. DEGs were declared for the interaction term as well for pair-wise comparisons according to the design ∼ CO2 + infection + CO2 : infection (absolute log2 FC ≥ 1, adjusted *P* ≤ 0.05, Wald test). GO term enrichment in biological processes (adjusted *P* ≤ 0.05, Fisher’s exact test) was performed in Panther (http://www.pantherdb.org/).

Metabolites were extracted from 3-10 mg of lyophilized plant material in 1 ml of 30% methanol (v/v) supplemented with 0.01% formic acid (v/v) (n = 6 independent biological replicates). Five microliters of metabolite extracts were injected and separated in a reverse Kinetex C18 analytical column (2.6 μm particle size, 50 mm × 2.1 m, Phenomenex) using a gradient of methanol and H2O supplemented with 0.01% formic acid. The flow rate was set to 0.3 ml/min. Samples were ionized in positive and negative ion modes for electrospray ionization (ESI) in a 40–1100 m/z range using an Acquity UPLC I-Class System interfaced to a hybrid quadrupole time-of-flight mass spectrometer, SYNAPT G2-S high-definition tandem mass spectrometry (MS/MS) detector (Waters, Milford, USA). Raw data were analysed depending on the ESI mode. Metabolome abundances were quantile normalised and fitted to linear mixed models to calculate significant differences for the interaction term as well for pair-wise comparisons using a post-hoc Dunnett’s test (adjusted *P* ≤ 0.05). Tentative annotation of metabolic features and KEGGs (adjusted *P* ≤ 0.05, Fisher’s exact test) were conducted with the MarVis-Suite 2.0 software (Kaever *et al*., 2015) with settings described in Garcia-Molina and Pastor (2014).

### Virus Induced Gene Silencing

VIGS assays were conducted using the tomato rattle virus (TRV) system as previously described by (Pastor-Fernández *et al*., 2024). Gene-specific fragments were selected using the Sol Genomics Network VIGS design tool (http://vigs.solgenomics.net/), amplified by PCR, and cloned into the pTRV2 vector after replacement of the PDS fragment. Target genes included two WRKY transcription factors (Solyc02g072190, Solyc02g093050) and five ERF transcription factors grouped into two independent silencing groups: ERF1 (Solyc09g089910, Solyc01g090340, Solyc08g078190) and ERF2 (Solyc04g078640, Solyc03g005520). Primer sequences and target regions are provided in **Table S17**.

For plant inoculation, Agrobacterium cultures carrying the corresponding pTRV2 constructs were pooled to obtain three VIGS combinations targeting WRKY, ERF1, or ERF2 genes, maintaining a final 1:1 ratio between pTRV1 and the combined pTRV2 cultures. Plants infiltrated with pTRV2_ev and pTRV2_PDS were used as empty vector and silencing controls, respectively. VIGS combinations were infiltrated into 3-week-old tomato plants, and *B. cinerea* infections were performed two weeks later.

### Bioinformatic analysis

For high-throughput analyses, quantile normalisation, data fitting to linear mixed models, hierarchical clustering for heatmaps and multiple comparisons were conducted with the RStudio packages *limma*, *pheatmap*, and *upSetR*, respectively. Gene Regulatory Networks (GRNs) were deconstructed using normalised counts for overlapping DEGs under infection between control and eCO2 conditions across the four genotypes (GRNb), adding specific DEGs under control conditions (GRNc), and DEGs due to the interaction factor, i.e., magnified under susceptible species (GRNs) using the *GENIE3* algorithm (Huynh-Thu *et al*., 2010). GRN topology was adjusted to a final density where node degree was stabilised according to the edge_testing function of *DIANE* (Cassan *et al*., 2021). Networks were visualised in Cytoscape (Shannon *et al*., 2003) and analysed using the RStudio package *igaph*. Enrichment in TF families was conducted according to the classification of TFs in PlantRegMap (https://doi.org/10.1093/nar/gkz1020) (adjusted *P* ≤ 0.05, Fisher’s exact test). Enrichments in putative cis-regulatory elements (pCREs) in promoter regions (1,000 bp upstream regions) of the target nodes in each GRN was computed by analysing the distribution of bona-fide identified motifs adapted from the Arabidopsis Gene Regulatory Information Server (https://agris-knowledgebase.org/AtcisDB/bindingsites.html; **Table S18**) compared to a 10,000 randomly selected promoters in the tomato transcriptome (adjusted *P* ≤ 0.05, Z-test).

## Data availability

Transcriptome and metabolome datasets have been deposited at the Gene Expression Omnibus (https://www.ncbi.nlm.nih.gov/geo/) and Metabolights (https://www.ebi.ac.uk/metabolights/).

## Author contributions

VP, RS-L and AG-M conceived the project, designed the experiments and supervised the progress, FB, MO-B and LY performed plant growth, infection assays, sample collection and analysis, VP and AG-M generated transcriptomic and metabolomic datasets. AG-M. conducted bioinformatic, network and statistical analyses. VP, RS-L and AG-M interpreted the results, prepared the figures and wrote the first draft of the manuscript. All authors revised and edited the manuscript. All authors read and approved the final version of the manuscript.

## Acknowledgements

We thank the Greenhouse Facility and Stress Program at CRAG for technical assistance to generate the biological material used in this study, and Dr. Cristian Vicent (Servei Central d’Instrumentació Científica, UJI) for his technical assistance with metabolomic analyses.

## Supplementary Material

**Figure S1.**
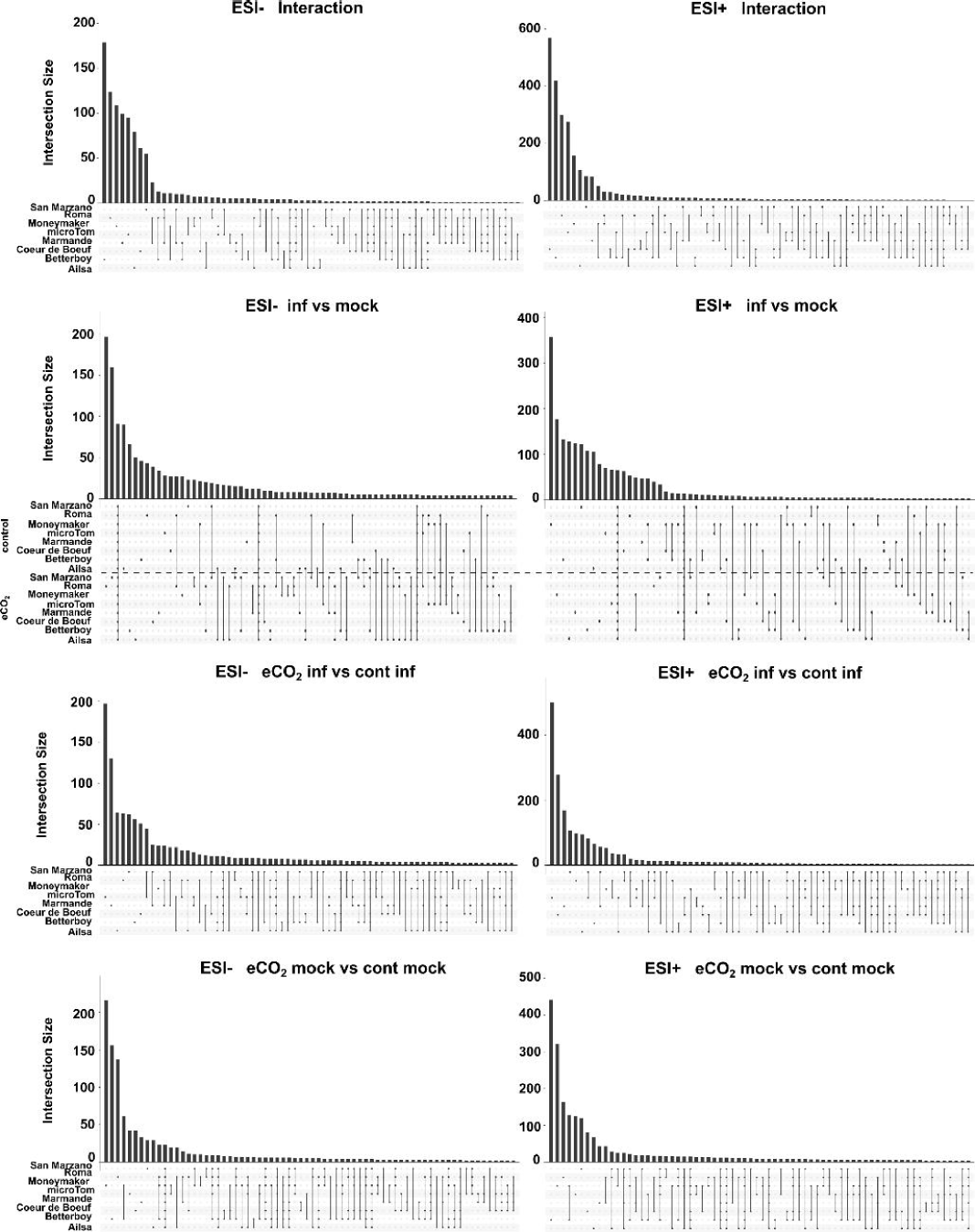
Multiple comparison of metabolome changes among varieties. Upset plots depict multiple comparisons of significantly accumulated metabolites (SAMs) for the indicated contrasts.

**Figure S2.**
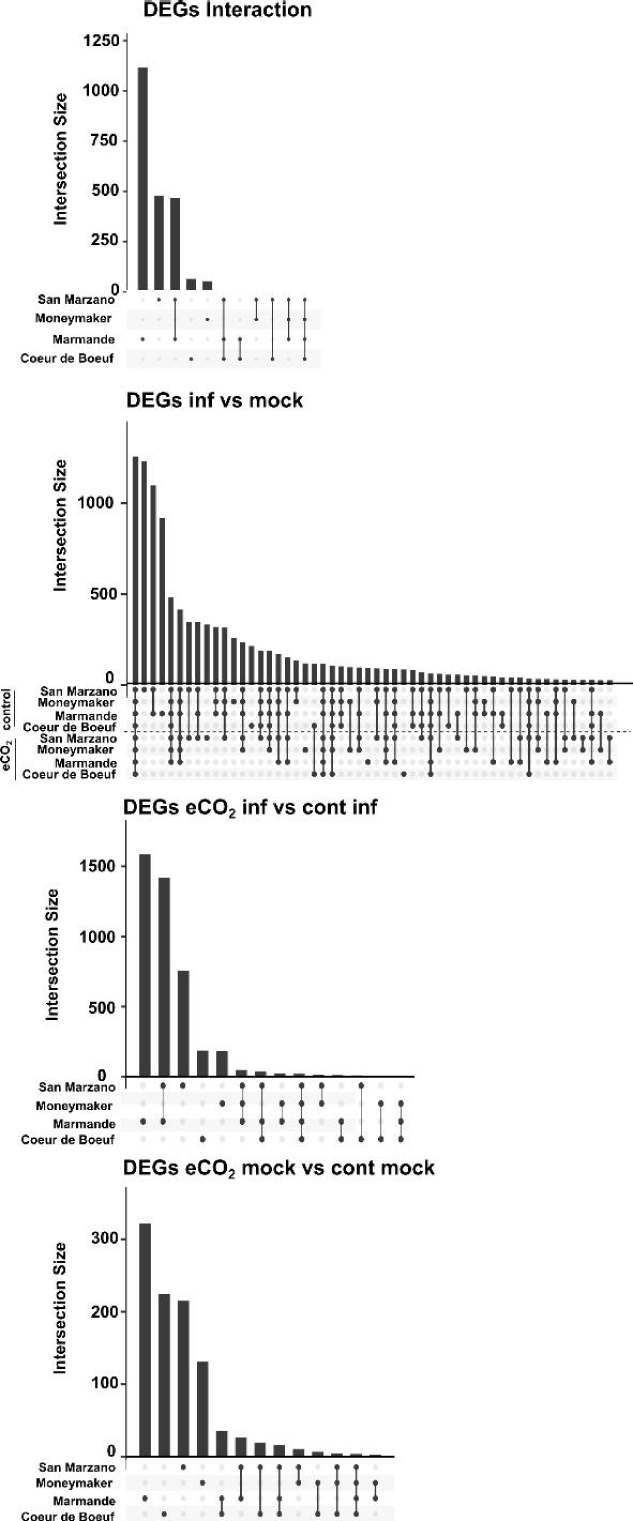
Multiple comparison of transcriptome changes among varieties. Upset plots depict multiple comparisons of differentially expressed genes (DEGs) for the indicated contrasts.

**Figure S3.**
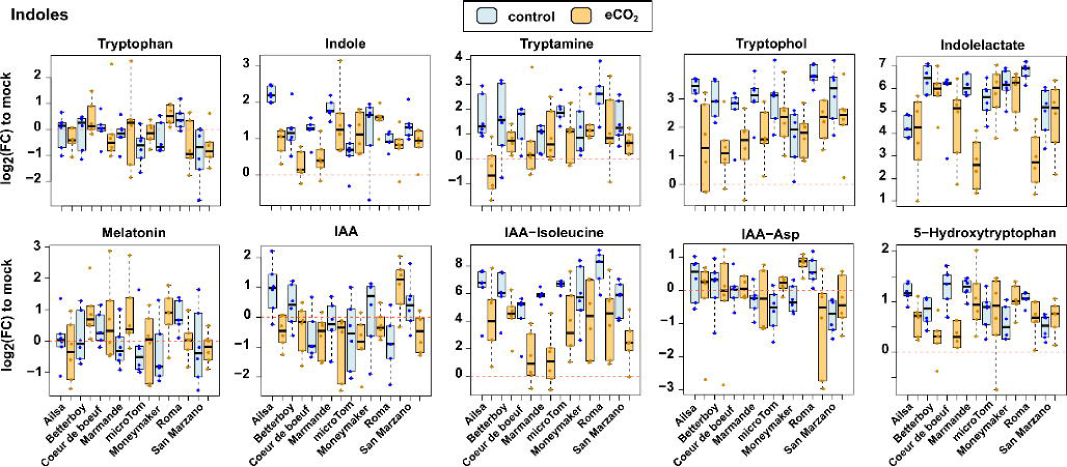
Reconfiguration of indole-derived metabolites by elevated CO₂ during *Botrytis cinerea* infection. Metabolome datasets were filtered for representative indole-related compounds. Boxplots depict log2-transformed fold-changes (FC) of infection-induced changes to mocks under control (light blue) and eCO₂ (light orange).

**Figure S4.**
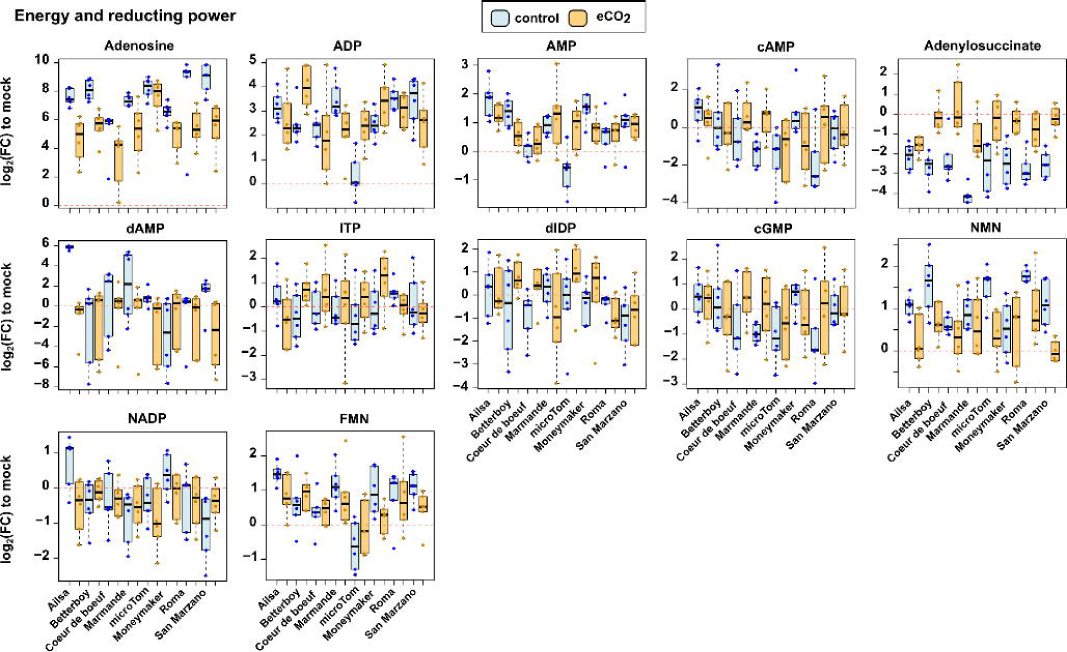
Reconfiguration of metabolites related to energy and redox state by elevated CO₂ during *Botrytis cinerea* infection. Metabolome datasets were filtered for energy and redox state -related compounds. Boxplots depict log2-transformed fold-changes (FC) of infection-induced changes to mocks under control (light blue) and eCO₂ (light orange).

**Figure S5.**
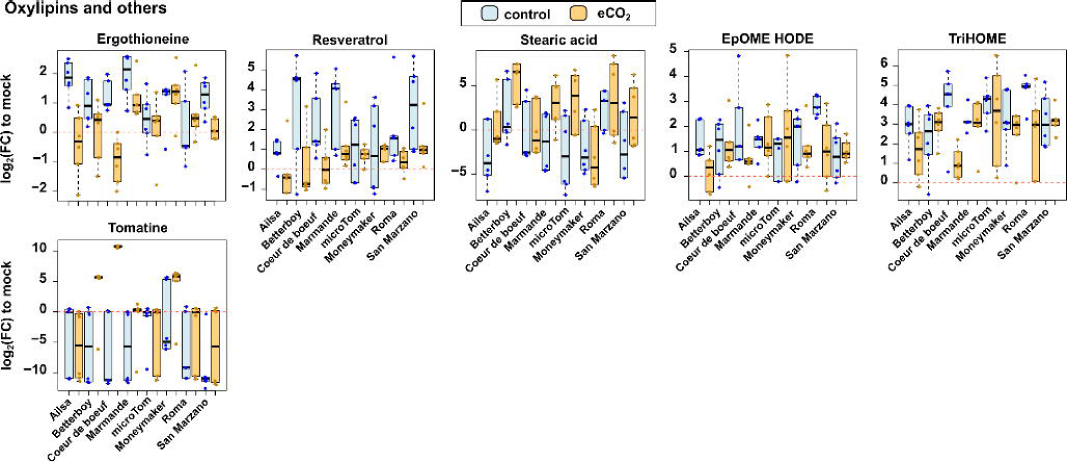
Reconfiguration of metabolites related to defensive compounds by elevated CO₂ during *Botrytis cinerea* infection. Metabolome datasets were filtered for defensive-related compounds. Boxplots depict log2- transformed fold-changes (FC) of infection-induced changes to mocks under control (light blue) and eCO₂ (light orange).

**Figure S6.**
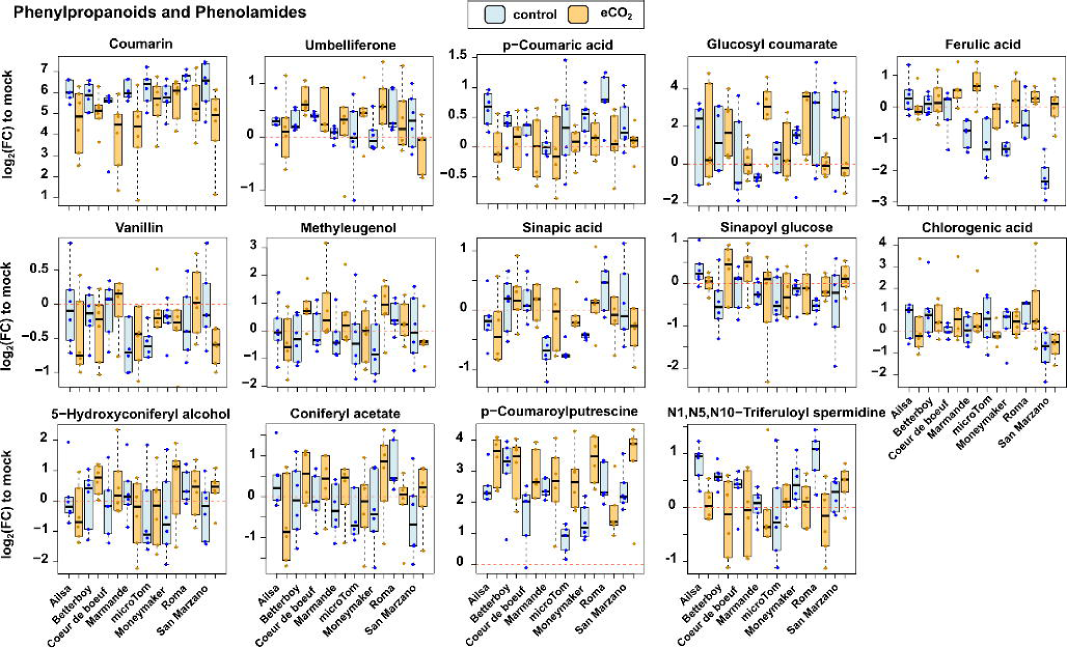
Reconfiguration of phenylpropanoids by elevated CO₂ during *Botrytis cinerea* infection. Metabolome datasets were filtered for phenylpropanoids. Boxplots depict log2-transformed fold-changes (FC) of infection-induced changes to mocks under control (light blue) and eCO₂ (light orange).

**Figure S7.**
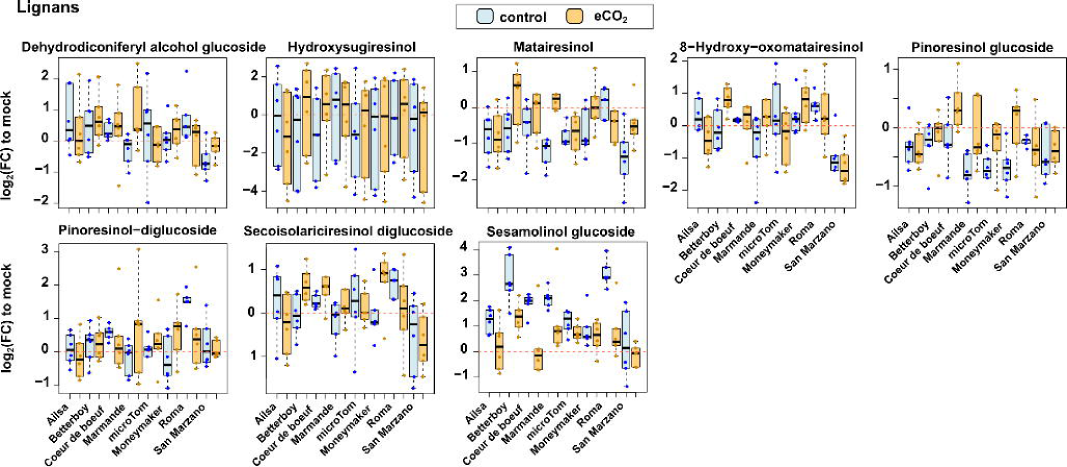
Reconfiguration of lignans by elevated CO₂ during *Botrytis cinerea* infection. Metabolome datasets were filtered for lignans. Boxplots depict log2-transformed fold-changes (FC) of infection-induced changes to mocks under control (light blue) and eCO₂ (light orange).

**Figure S8.**
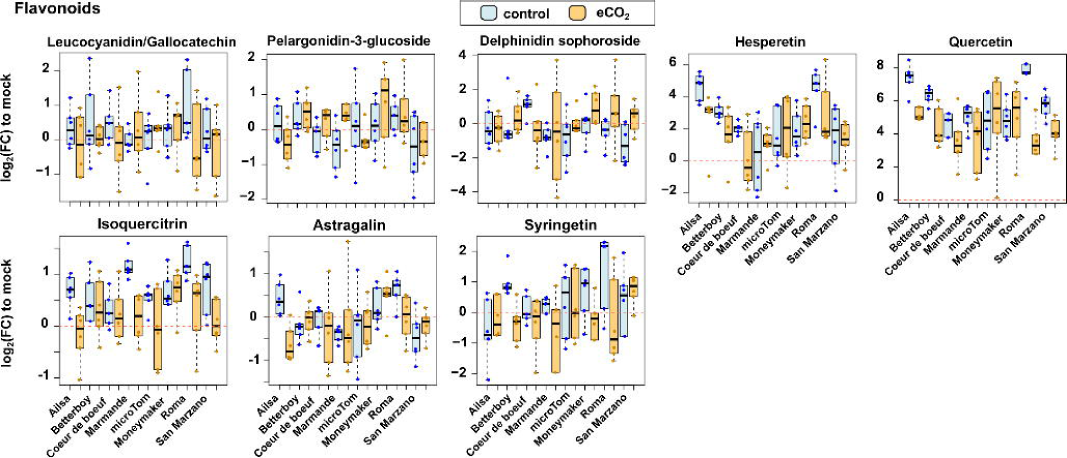
Reconfiguration of flavonoids by elevated CO₂ during *Botrytis cinerea* infection. Metabolome datasets were filtered for flavonoids. Boxplots depict log2-transformed fold-changes (FC) of infection-induced changes to mocks under control (light blue) and eCO₂ (light orange).

**Table S1. Lists of compounds detected in the metabolome profiling. Table S2. Metabolome changes due to infection and CO2 environments.**

**Table S3. Tentative annotation of metabolites changing as result of the inteaction between infection and CO2 conditions.**

**TableS4. Functional annotation of significantly altered metabolites with consistent patterns due to the interaction of the infection and the environmental CO2.**

**Table S5. Lists of differentially expressed genes due to infection and environmental CO2.**

**Table S6. Classification of trajectories of differentially expressed genes due to the infection and environmental CO2.**

**Table S7. Functional annotation of differential transcripts due to the interaction between infection and environmental CO2 depending on the expression pattern.**

**Table S8. Shared infection-responsive transcripts.**

**Table S9. Functional annotation of infection-responsive transcripts under control conditions. Table S10. Gene regulatory networks.**

**Table S11. Enrichment in transcription factor families in the Gene Regulatory Networks. Table S12. Functional annotation of targets in each Gene Regulatory Network.**

**Table S13. Central transcription factors in Gene Regulatory Networks.**

**Table S14. Targets of central transcription factors in the Gene Regulatory Networks.**

**Table S15. Functional annotation of nodes in the regulatory module of the Gene Regulatory Networks. Table S16. Targeted metabolome analysis.**

**Table S17. Oligonucleotides used in this study. Table S18. List of putative cis-regulatory elements.**

